## Supplementary Information for "High resolution species assignment of *Anopheles* mosquitoes using *k*-mer distances on targeted sequences"

#### Section 1: Species labels

The wild-caught mosquitoes in the reference database were labelled either morphologically or confirmed with species diagnostic PCR. Species identities were further determined by Sanger sequencing the molecular barcodes Cytochrome *c* oxidase I (COI) and Internal transcribed spacer 2 (ITS2). Both markers were compared to the NCBI database to assess species identity, and COI was also compared to the BOLD database. (See (Makunin et al. 2022) for more information).

Some partner labels were revised when distances between the samples in the reference dataset suggested mislabelling. Preferably the relabelling was further supported by molecular barcodes. Relabelled samples are:

- Amar-5: *An. marshallii → An. gambiae*. The average distance to samples in the *An. marshallii* complex is 0.54 (0.52-0.58), while the average distance to samples in *An. gambiae/coluzzii* is 0.09 (0.07-0.13). Additionally, both COI and ITS2 have *An. gambiae* as best hit.
- Amar-42: *An. marshallii → An. jebudensis*. The average distance to samples in the *An. marshallii* complex is 0.38 (0.35-0.42), while the distance to *An. jebudensis* is 0.05. Unfortunately, neither *An. marshallii* nor *An. jebudensis* is present in the barcode databases.
- Amar-3-1: *An. marshallii → An_marshallii_cp_sp1*. It has the lowest distances to samples of *An. hancocki, An. brohieri, An. demeilloni, An. theileri*, on average 0.26 (0.22-0.29), while the distances of *An. hancocki, An. brohieri* and *An. demeilloni* samples to each other is on average 0.03 (0.02-0.04) and the distances of the *An. theileri* samples is 0.07 (0.05-0.08). The average distance between the group containing *An. hancocki, An. brohieri* and *An. demeilloni* and the *An. theileri* is 0.29 (0.25-0.33). So it seems like *Amar-3-1* does not belong to either of these two groups, but it is as related to these groups as they are to each other. Unfortunately it does not have any informative matches on the molecular barcodes. Therefore this sample is labelled as an unnamed species in the *An. marshallii* complex.
- Adem-15: *An. demeilloni → Myzomyia_sp1*. Its lowest distance to another sample in the database is 0.39, to an *An. funestus*. This is a much higher distance than a sample usually has to other samples of the same or of a closely related species. In particular, the distance of *An. hancocki, An. brohieri* and *An. demeilloni* samples to each other is 0.03 (0.02-0.04), while the average distance of Adem-15 to these samples is 0.41 (0.39-0.44). As it does not show similarity to any of the samples in the database, but it is closer to almost all samples in the *Myzomyia* series than to almost all samples outside the *Myzomyia* series, it is labelled as an unnamed species in the *Myzomyia* series.
- Acol-645: *An. coluzzii → An_gambiae_cp_sp1*. This sample is different from other *An. coluzzii* and *An. gambiae*, this was found both in the sample-pair differences and in the subsequent VAE method which is tailored at the *An. gambiae* complex. The molecular barcodes also suggest that this sample is not *An. coluzzii* or *An. gambiae*,

but that it is a member of the complex. Therefore it is labelled as an unnamed species in the *An. gambiae* complex.

- Apal-257: *An. paludis → An_coustani_cp_cl3*. Its smallest distances are to samples of *An. tenebrosus, An. ziemanni, An. coustani* and *An. paludis*, on average 0.29 (0.28-0.32). We found that the samples of these species split into two clades and the distance of Apal-257 to either of these clades was much higher than the distances between samples in the same clade. Additionally, the average distances between the two clades is 0.18 (0.14-0.21), so Apal-257 is also further away from the two clades than they are from each other. Yet, it is closer to these two clades than it is to *An. sinensis* and *An. hyrcanus*, which are the next closest species. Regarding the molecular barcodes, Apal-257 matches to *An. coustani* on COI, like most samples in the *An. coustani* complex. On ITS2 it matches to *An. junlianensis* and *An. yatsushiroensis*, species in the hyrcanus group. We therefore decided to relabel it to a third clade in the *An. coustani* group.
- *An. nili* samples. Anils-7: *An. nili s.s. → An_nili_gp_sp1*. Anil-237 & Anil-239: *An. nili → An_nili_gp_sp2.* Anil-233, Anil-236 & Anil-238: *An. nili → An_nili_gp_sp3.* Both the distances and alignment of the molecular barcodes support this split. It is possible that some of these samples represent different member species of the *An. nili* group, which have been shown from cytogenetic analysis to differ substantially (Sharakhova et al. 2013), but molecular barcodes are not yet available in public databases for all member species.

| dist | *An_nili_gp_sp2* | *An_nili_gp_sp3* |
|---|---|---|
| *An_nili_gp_sp1* | 0.19 (0.19-0.20) | 0.24 (0.23-0.26) |
| *An_nili_gp_sp2* | 0.02 | 0.24 (0.24-0.26) |
| *An_nili_gp_sp3* | | 0.04 (0.03-0.05) |

- *An. hyrcanus* samples. VBS00085 & VBS00086: *An. hyrcanus → An_hyrcanus_gp_sp1*. VBS00082 & VBS00083: *An. hyrcanus → An_hyrcanus_gp_sp2.* The distances within these pairs are 0.08 for both pairs. The distances between the pairs are 0.35 (0.34-0.36). The molecular barcode matches support the split. Even though hyrcanus is present in all databases, there are few matches to it. On COI in both BOLD and NCBI, the first pair matches to *nitidus* and the second pair to *crawfordi*; these species are in two distinct subgroups of the hyrcanus group. On ITS2 the first pair matches to *hyrcanus* and then *nitidus*, and the second pair matches to *sinensis* (*sinensis* is in the main *hyrcanus* group, not in either of the aforementioned subgroups). The alignments of COI and ITS2 sequences for these four samples also clearly support the split into pairs.
- Samples in the *An. coustani* complex. This complex contains the species *An. tenebrosus, An. ziemanni, An. coustani* and *An. paludis*. The between sample distances indicate a split into two clades (disregarding sample Apal-257, discussed above), however, the split is not correlated with the species labels. The proposed clades are *An_coustani_cp_cl1* containing Aten-191, Aten-185, Azie-334, Acou-956, Acou-959, Acou-962 and *An_coustani_cp_cl2* containing Aten-333, Aten-79, Aten-954, Azie-1032, Azie-1055, Azie-70, Azie-77, Acou-71, Acou-80, Apal-81. The average distances between members of the same clade is 0.07 (0.05-0.09) and 0.07 (0.05-0.10) respectively, while the distance between members of different clades is 0.18 (0.14-0.21). The barcode matches are to *An. coustani* for the vast majority of the

samples, even though COI sequences for *An. ziemanni* and *An. tenebrosus* are present in both BOLD and NCBI database. Alignment of the COI sequences shows extremely little variation. Alignment of ITS2 does support a split into the two proposed clades.

All labels and groupings are displayed in Supplementary Table ST1. In the majority of cases, the fine species-groups are supported by at least one molecular marker and not contradicted by different species labels. Some species are not present in one or both databases, so for these a match to a closely related species is allowed. The exceptions are:

- *An. jebudensis*: one sample was labelled as *An. jebudensis*, the other one originally as *An. marshallii*, but showed sufficient evidence for relabelling. Neither *An. jebudensis* nor *An. marshallii* is present in either database. On ITS2 they both match to *An. moucheti*, which is the closest species in the dataset, neither has a match in BOLD, and on COI in the NCBI database the best matches are to *An. lindesayi* and *An. sawyeri*, which are in the *Anopheles* and *Nyssorhynchus* subgenus respectively. However, it has to be noted that the NCBI database does not contain an *An. moucheti* COI sequence and the best hits are based on 81% and 91% identity respectively.
- *An. rhodesiensis*: All samples match to uninformative ‘Anopheles_sp’; even though a COI sequence is available in BOLD and NCBI. The best hit to a named species in the dataset is to *An. funestus* for COI and *An. aconitus* for ITS2.
- *An. jamesii*: Is predicted for BOLD and ITS2, but not for COI in the NCBI database, even though an *An. jamesii* COI sequence is present there.
- *An. maculatus A*: has a few matches to *An. sawadwongporni*, which is also a member of the maculatus group, even though sequence of *An. maculatus A* is present in all databases.
- *An. rampae*: the four different samples match to *An. rampae(2x), An. sawadwongporni(1x)* and *An. maculatus(1x)* in the BOLD database; all these are in the same species group. In the NCBI database, all COI hits are for *An. maculatus* (*An. rampae* COI not present in database), and all ITS2 hits are *An. rampae* and the uninformative *Anopheles_sp.*
- *An. balabacensis*: matches to *An. introlatus* on COI in both databases, even though *An. balabacensis* sequence is present. The two species are in the same complex.
- *An. carnevalei*: all samples match to *An. carnevalei* on BOLD and ITS2. *An. carnevalei* sequence is not present in NCBI database for COI; 4 out of 5 samples match to *An. nili,* one to *An. darlingi*, which is in a different subgenus.
- *An. dureni*: sequence not present in either database; on ITS2 one sample matches to *An. minimus* and one to *An. leesoni*, which are both in different groups; but placement of *An. dureni* is not very clear on the species tree.
- *An. vinckei*: sequence not present in either database; on COI there are distant matches to *An. gambiae* and *An. maculatus*, which are both in different groups; but placement of *An. vinckei* is not very clear on the species tree (seems to be close to *An. dureni*).
- *An. coustani* group: we have samples of four species, *An. coustani* (present in all databases)*, An. paludis* (present in no databases)*, An. ziemanni* (COI present in both databases) and *An. tenebrosus* (COI present in both databases). In BOLD, all samples match to *An. coustani*. In the NCBI database, on COI most matches are to *An. coustani*, there is one match to *An. ziemanni*, and a few matches to

*Anopheles_cf.* On ITS2 there are many matches to *Anopheles_sp.* and *Anopheles_cf.*, and the remaining matches are to *An. coustani* and one sample to *An. junlianensis* and *An. yatsushiroensis* in the hyrcanus group (this is the *An. paludis* sample, that is further removed from all other samples in this complex). These samples also show a 'checkerboard' pattern, which splits them into two clades, which are not correlated with species labels.
- *An. barbirostris*: there are matches to *An. barbirostris* and to the closely related *An. dissidens*. However, the samples do not form two separate groups based on the distances.
- *An. oryzalimnetes*: no Sanger sequencing done.
- *An. cruzii:* no Sanger sequencing done.
- *An. bellator*: no Sanger sequencing done.

Lastly, there are samples which are further than 0.10 distance away from other samples in the reference database representing the same species. Above we have discussed some samples where there was good evidence to adjust the species label, but there are also some for which there is good reason to retain the partner label. Those are:
- VBS00001: *An. annularis*. It is a bit more diverged from the other three *An. annularis* samples, which causes it to fall just above the threshold. However, the distances are not very different from the distance between the other *An. annularis*, the next closest sample in the reference set are much further away and the molecular barcodes support the partner label.
- Agam-37: *An. gambiae*. It is a bit more diverged from the other *An. gambiae* that were sequenced by the panel, which causes it to fall just above the threshold. But is it closer to *An. gambiae* than to other samples in the reference dataset. The molecular barcodes also support the partner label.
- VBS00149 & VBS00150: *An. tessellatus*. They are 0.11 distance from each other, but for both the molecular barcodes match to *An. tessellatus* and the next closest sample in the database is at 0.44 distance.
- anopheles-sinensis-chinascaffoldsasinc2 & anopheles-sinensis-sinensisscaffoldsasinc2: *An. sinensis*. They are 0.11 distance from each other and that is still clearly closer than the next best match in their sister species *An. hyrcanus* (namely *An_hyrcanus_gp_sp2*).

### Section 2: Thresholds defining species-groups

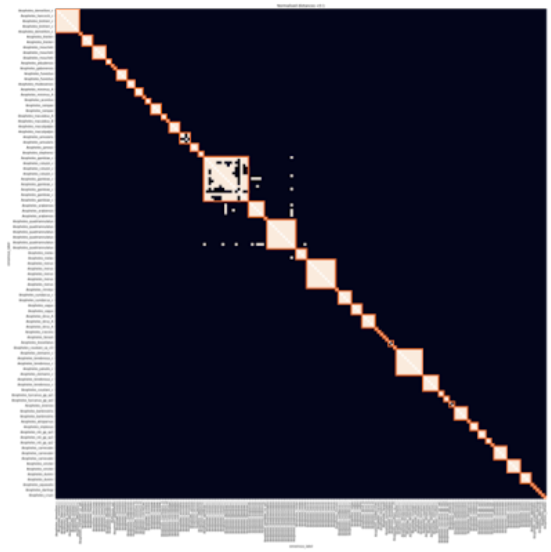

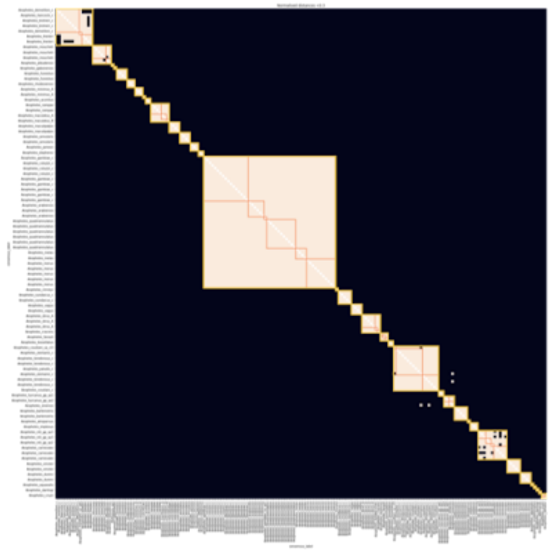

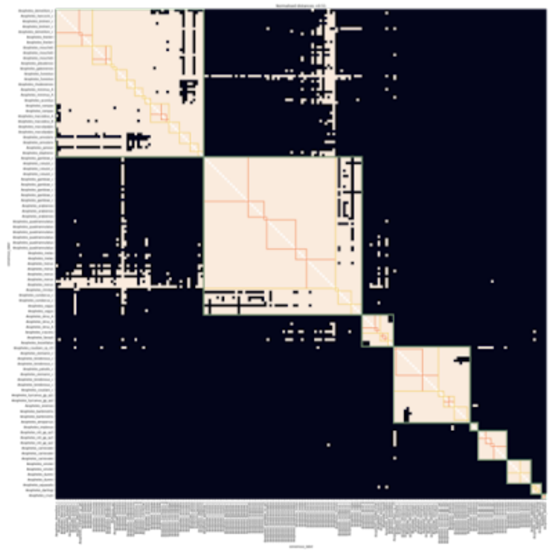

**Supplementary Figure 1:** Thresholds used to define species-groups. From top to bottom, the thresholds are 0.1, 0.3 and 0.51 and they are used to define the fine, intermediate, and coarse level species groups, respectively. The samples from the reference database are along the x- and y-axis in the same order as in Figure 1. The entries in the heatmap are coloured peach if the 8-mer distance between the corresponding samples is less than the threshold and black if it is greater or equal the threshold. The orange squares in all three panels correspond to fine level species-groups, the yellow squares in the middle and lower panel to intermediate level species-groups and the olive squares in the lower panel to coarse level species-groups.

### Section 3: Species-groups assignment results reference database

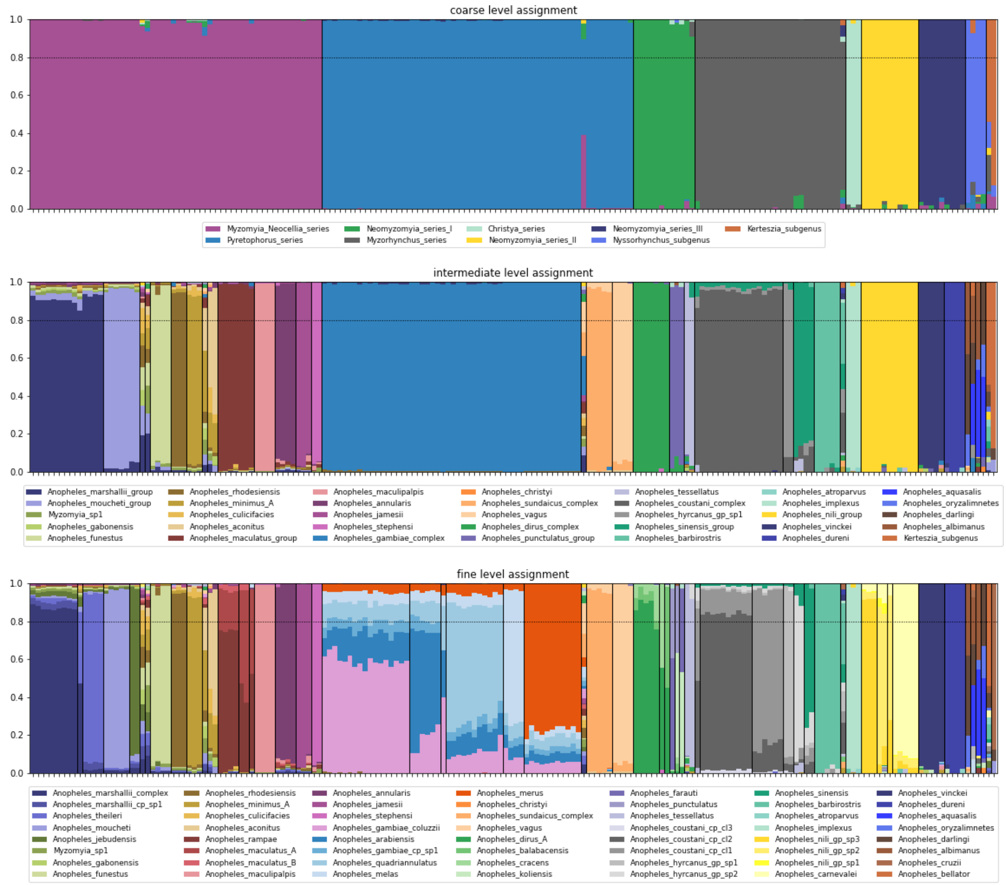


**Supplementary Figure 2:** Species-group assignment proportions. From top to bottom at the coarse, intermediate and fine level. Samples are ordered along the x-axis in the same order as the tree in figure 3. For each sample, the overall assignment proportions are plotted as a bar, with colours indicating the species-groups. A sample has to have an assignment proportion of at least 0.8 for a certain species-group to be classified as a member of that species-group, else it remains unassigned. Vertical bars separate the different species-groups. The horizontal bar represents the 0.8 assignment threshold (but note that the assignment proportions are plotted in the same order for every sample, not from largest to smallest).

### Section 4: Parameter choices for the VAE

Our VAE architecture is inspired by popVAE (Battey, Coffing, and Kern 2021). We used their default parameter choices for learning rate, training iterations and validation proportion as well as the ‘elu’ propagation between the layers of the neural network and the ‘linear’ activation function to transform the output from the encoder to the latent space variables. Because we model *k*-mer counts as independent Poisson variables, we use the ‘softplus’ activation function to transform the output from the last layer of the decoder.

To set the width and depth of the encoder and decoder, we ran a grid search over a combination of widths and depths. We kept the width and depth the same for the decoder and the encoder and we trained the VAE with three different seeds for each width-depth combination. For high width values (more than 500 nodes) the run time required to train the VAE was very long and the process would often crash when conducted on a desktop. For very small networks (width 32, depth 4) the resolution was poor, but in the middle range the visible structure was not severely affected by the choice of width and depth, so we went with width 128 and depth 6, which gave a good resolution and reasonable run time.

As mentioned in the main text, the loss function contains a parameter, *w*, that controls the relative importance of the data driven term and the regularisation term. We ran a grid search ranging over four orders of magnitude, training with three different seeds for each search point. For small values of *w*, which gives higher weight to regularisation term, the resulting latent space dimension shows a strong correlation between the latent space dimensions, suggesting that only highly differentiated samples can overpower the effect from the regularisation term. For high values of *w*, the training process of the VAE becomes more unstable: it crashes sometimes and the resulting latent space projections look less similar to those obtained with the same *w* value, but a different seed. We chose *w* equal to 1000, which resulted in reasonably stable projections, with good visible structure.

To determine the number of latent space dimensions, we ran a grid search over 2, 3 and 4 latent space dimensions. In fact, the grid search for *w* and the number of latent space dimensions was run simultaneously, but there seemed to be little interaction between *w* and the number of latent space dimensions. For the training set containing only *An. gambiae*, *An. coluzzii* and *An. arabiensis*, adding a third or fourth dimension did not add to the visible structure, so in this case we opted for two latent space dimensions for visualisation purposes. However, for the full training set GCref v1, in the two-dimensional projection, several species clusters are very close to each other, making this hard to use for species assignment. Adding a third dimension resolves this problem, even though the second and third dimensions are strongly correlated (see Figure 4 in the main text). The benefit of the strong correlation is that we can visualise the structure in two dimensions.

### Section 5: Geographic stability

Here we test the robustness of our VAE projection by removing all samples collected in one geographic location from the training set and then projecting those samples and the validation set and comparing these results to those obtained using the VAE trained on the full training set. In particular, we will pay attention to the separation of the species clusters, the classification accuracy of the validation set and the visible structure within the species clusters.

For this analysis, we use a subset of the reference dataset GCref v1 and the validation set GCval v1, containing only the species *An. coluzzii*, *An. gambiae* and *An. arabiensis*, because for the other species in the *An. gambiae* complex we have very few samples, the samples in this dataset are collected at only one or two locations and the species have more limited geographic ranges than *An. coluzzii*, *An. gambiae* and *An. arabiensis*. With only these species, two latent space dimensions are sufficient to exhibit the relevant structure, so that is what we will use here. We have performed these drop-outs for a subset of the geographic locations, selecting for those we thought would be most likely to affect the structure of the projection.

Below, we discuss the drop-out experiments in more detail, but to summarise: overall, the separation of the species clusters remained intact and the ability to classify species of the samples in the training set and and in the validation set was not significantly affected. Samples from The Gambia and Guinea-Bissau could be less accurately classified when no samples from this geographic region were included in training the VAE, but for all other locations we tested, the ability to classify species remained the same. The degree to which geographic structure is visible within the species clusters was somewhat affected for certain collection countries. However, the visible structure never completely disappeared, nor did we see a considerably different geographic structure, it was merely that the structure became a bit more blurred or more condensed than in the original projection.

#### Angola

We removed 10 *An. coluzzii* samples from the training set, see Supplementary Figure 1. The separability of species clusters is similar compared to the projection of the VAE trained on the full dataset. The geographic structure within the *An. gambiae* and *An. arabiensis* clusters remains intact, but the *An. coluzzii* cluster loses some of its visible structure. Angola is on the edge of the geographical range of the *An. coluzzii* samples in this dataset and these samples are also on the very top of the VAE projection, suggesting that they are responsible for a considerable amount of the structure within *An. coluzzii*.

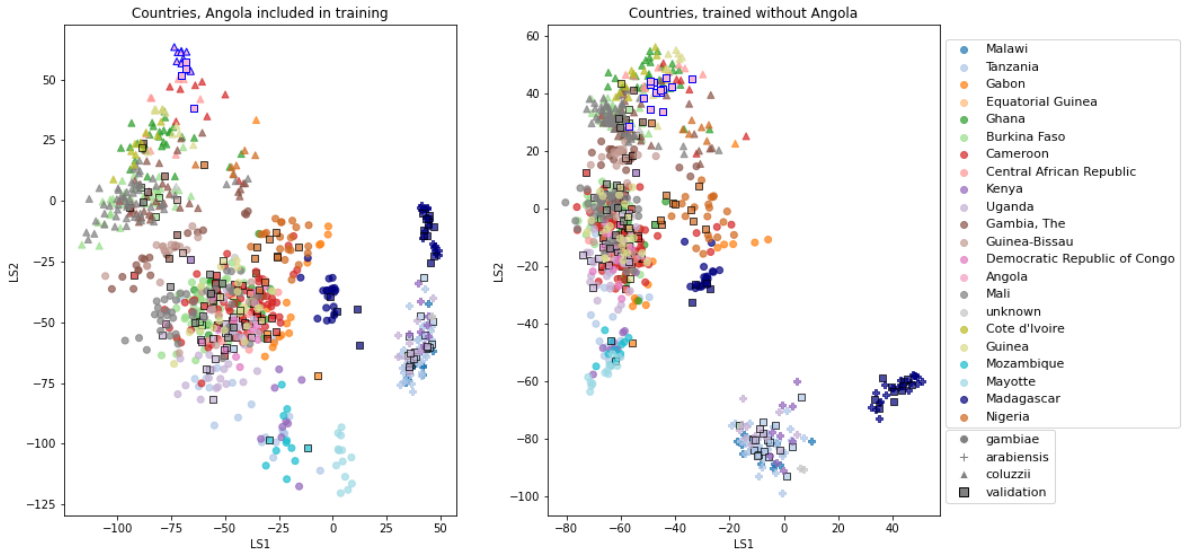


**Supplementary Figure 3:** VAE projections of *An. arabiensis*, *An. coluzzii* and *An. gambiae* from different geographic locations. **Left:** samples from Angola included in VAE training. **Right:** samples from Angola excluded from VAE training. Samples are coloured by country of collection. Squares are validation samples (not used in VAE training), triangles are *An. coluzzii* individuals, circles are *An. gambiae* individuals and crosses are *An.*

*arabiensis* individuals. Samples from Angola are highlighted with a blue edge, all other validation samples have a black edge.

##### Cameroon

We removed 10 *An. coluzzii* and 66 *An. gambiae* from the training set, see Supplementary Figure 2. The separability of the species clusters is similar compared to the projection of the VAE trained on the full dataset. The geographic structure within the *An. coluzzii* and *An. arabiensis* clusters remains intact, but the *An. gambiae* cluster loses some visible structure in its main subcluster, while the structure in the other four subclusters (formed by samples from The Gambia and Guinea-Bissau; by samples from Nigeria and Gabon; by samples from Madagascar and by samples from various East African countries respectively) remains largely intact. This is probably due to the fact that the samples from Cameroon form a large proportion of the samples in the main *An. gambiae* cluster.

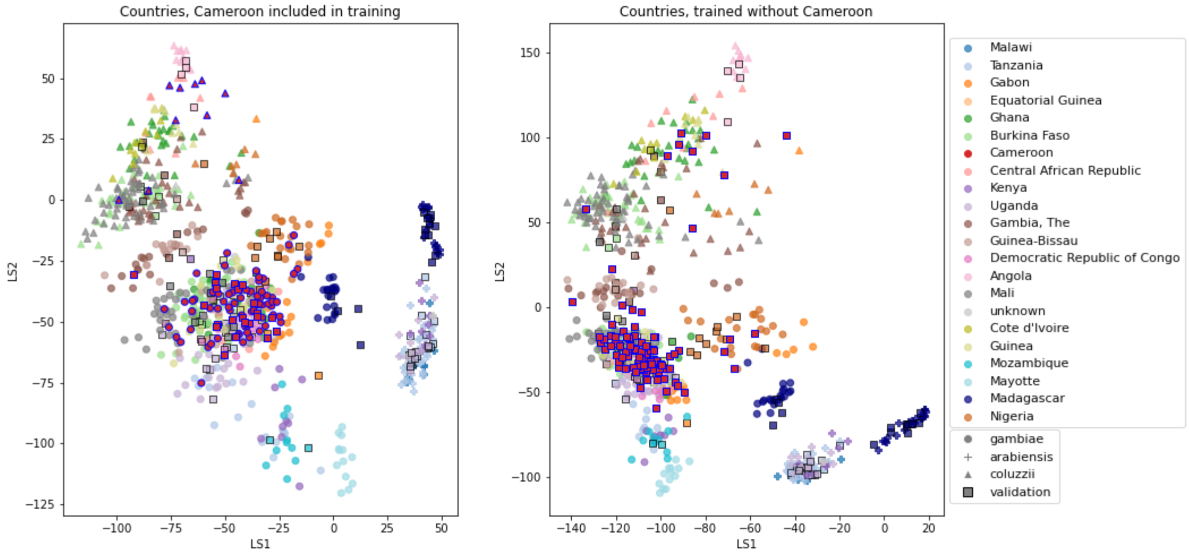


**Supplementary Figure 4:** VAE projections of *An. arabiensis*, *An. coluzzii* and *An. gambiae* from different geographic locations. **Left:** samples from Cameroon included in VAE training. **Right:** samples from Cameroon excluded from VAE training. Samples are coloured by country of collection. Squares are validation samples (not used in VAE training), triangles are *An. coluzzii* individuals, circles are *An. gambiae* individuals and crosses are *An. arabiensis* individuals. Samples from Cameroon are highlighted with a blue edge, all other validation samples have a black edge.

##### The Gambia and Guinea-Bissau

We removed 33 *An. coluzzii* and 19 *An. gambiae* from The Gambia and 24 *An. gambia* from Guinea-Bissau from the training set, see Supplementary Figure 3. Considering the training samples, the separability of the species clusters is very clear. However, the classification accuracy of the projected samples from The Gambia and Guinea-Bissau is only 70%. This is probably due to the fact that these samples form the border between the *An. coluzzii* and *An. gambiae* clusters in the projection of the VAE trained on the complete dataset. So by leaving those samples out, we miss the important label information necessary to classify the edge cases. The subclusters of the *An. gambiae* cluster are arguably more pronounced than in the original projection, but the structure within the subclusters of the *An. gambiae* cluster and in the entire *An. coluzii* cluster reduces.

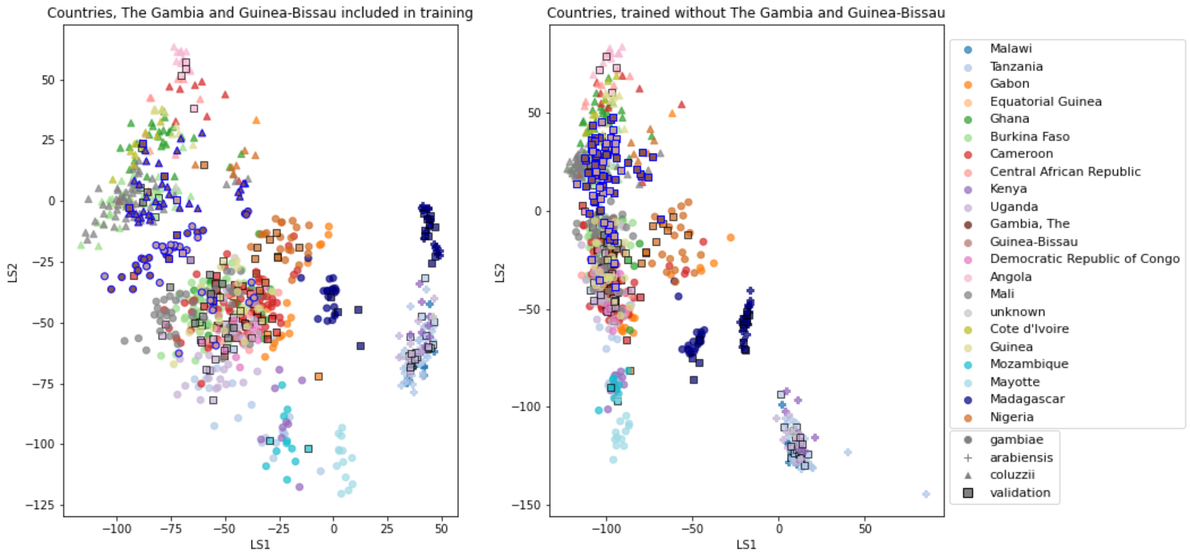


**Supplementary Figure 5:** VAE projections of *An. arabiensis*, *An. coluzzii* and *An. gambiae* from different geographic locations. **Left:** samples from The Gambia and Guinea-Bissau included in VAE training. **Right:** samples from The Gambia and Guinea-Bissau excluded from VAE training. Samples are coloured by country of collection. Squares are validation samples (not used in VAE training), triangles are *An. coluzzii* individuals, circles are *An. gambiae* individuals and crosses are *An. arabiensis* individuals. Samples from The Gambia and Guinea-Bissau are highlighted with a blue edge, all other validation samples have a black edge.

#### Madagascar

We removed 20 *An. arabiensis* and 18 *An. gambiae* from the training set, see Supplementary Figure 4. The separability of the species clusters is similar compared to the projection of the VAE trained on the full dataset and the geographic structure within the three species clusters remains largely intact. When the Madagascar samples are included in training the VAE, they form a tight subcluster, both for *An. gambiae* and *An. arabiensis*. When they are not included in training, these clusters stand out less, due to the fact that the VAE does not recognise the features that make them stand out.

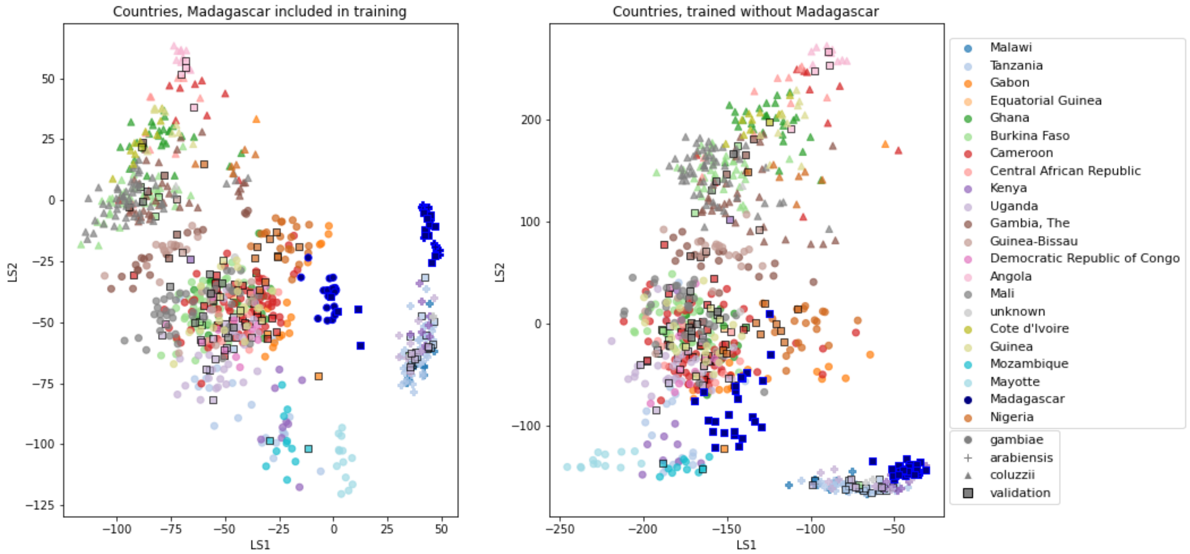


**Supplementary Figure 6:** VAE projections of *An. arabiensis*, *An. coluzzii* and *An. gambiae* from different geographic locations. **Left:** samples from Madagascar included in VAE training. **Right:** samples from Madagascar excluded from VAE training. Samples are coloured by country of collection. Squares are validation samples (not used in VAE training), triangles are *An. coluzzii* individuals, circles are *An. gambiae* individuals and crosses are *An. arabiensis* individuals. Samples from Madagascar are highlighted with a blue edge, all other validation samples have a black edge.

#### Mali

We removed 57 *An. coluzzii* and 43 *An. gambiae* from the training set, see Supplementary Figure 5. The separability of the species clusters is similar compared to the projection of the VAE trained on the full dataset. The visible geographic structure within the three species clusters reduces a bit. This could be because we removed such a large number of samples.

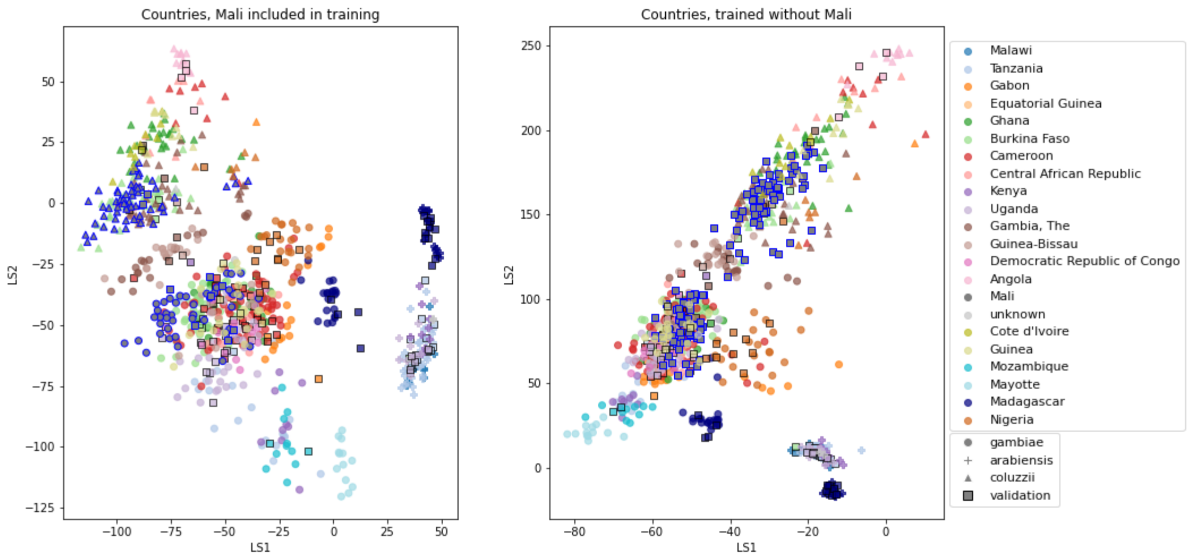


**Supplementary Figure 7:** VAE projections of *An. arabiensis*, *An. coluzzii* and *An. gambiae* from different geographic locations. **Left:** samples from Mali included in VAE training. **Right:** samples from Mali excluded from VAE training. Samples are coloured by country of collection. Squares are validation samples (not used in VAE training), triangles are *An. coluzzii* individuals, circles are *An. gambiae* individuals and crosses are *An. arabiensis* individuals. Samples from Mali are highlighted with a blue edge, all other validation samples have a black edge.

#### Nigeria

We removed 6 *An. coluzzii* and 20 *An. gambiae* from the training set, see Supplementary Figure 6. The separability of the species clusters is similar compared to the projection of the VAE trained on the full dataset and the geographic structure within the three species clusters remains largely intact. When the Nigerian samples are included in training the VAE, the *An. gambiae* individuals are more separated from the main *An. gambiae* cluster than when they are projected using the VAE excluding them from training. This is because the VAE has not been trained to recognise the features that distinguish these Nigerian *An. gambiae* individuals from the other samples.

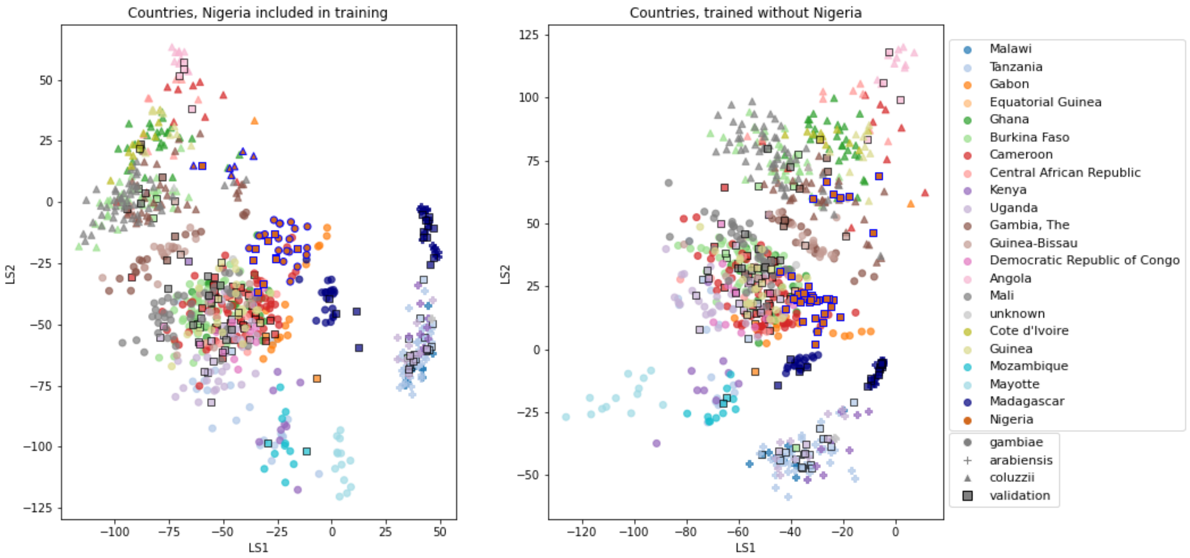


**Supplementary Figure 8:** VAE projections of *An. arabiensis*, *An. coluzzii* and *An. gambiae* from different geographic locations. **Left:** samples from Nigeria included in VAE training. **Right:** samples from Nigeria excluded from VAE training. Samples are coloured by country of collection. Squares are validation samples (not used in VAE training), triangles are *An. coluzzii* individuals, circles are *An. gambiae* individuals and crosses are *An. arabiensis* individuals. Samples from Nigeria are highlighted with a blue edge, all other validation samples have a black edge.

#### Tanzania

We removed 37 *An. arabiensis* and 20 *An. gambiae* from the training set, see Supplementary Figure 7. The separability of the species clusters is similar compared to the projection of the VAE trained on the full dataset. The subclusters of *An. gambiae* and *An. arabiensis* remain well separated, but the visible geographic structure within the (sub)clusters reduces a bit.

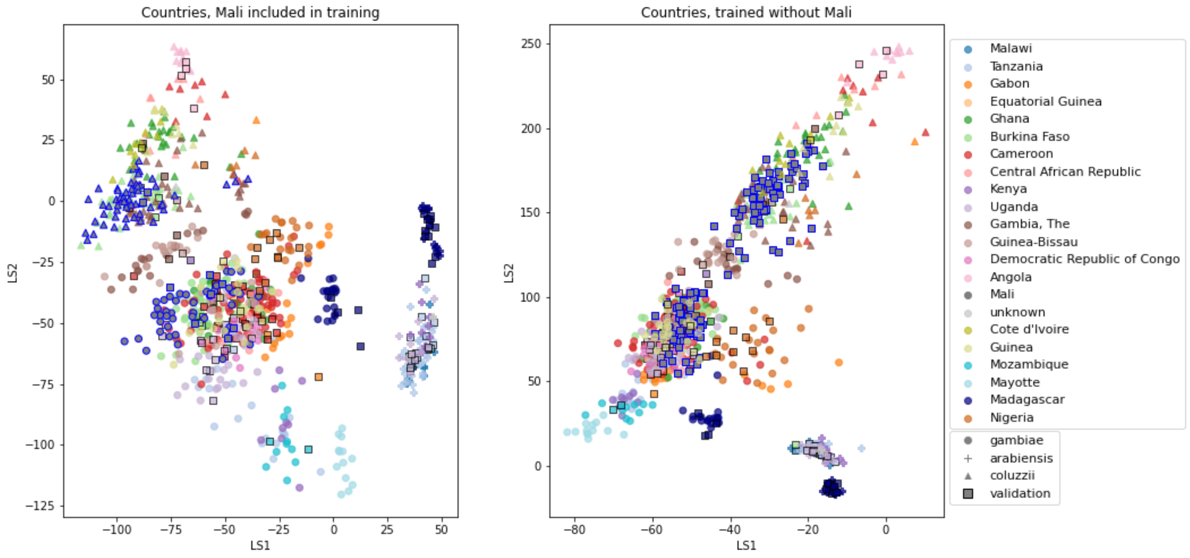


**Supplementary Figure 9:** VAE projections of *An. arabiensis*, *An. coluzzii* and *An. gambiae* from different geographic locations. **Left:** samples from Tanzania included in VAE training. **Right:** samples from Tanzania excluded from VAE training. Samples are coloured by country of collection. Squares are validation samples (not used in VAE training), triangles are *An. coluzzii* individuals, circles are *An. gambiae* individuals and crosses are *An. arabiensis* individuals. Samples from Tanzania are highlighted with a blue edge, all other validation samples have a black edge.
